# The road to nowhere: geolocation-by-genotype traces large-scale yellow-browed warbler vagrancy to central Siberia

**DOI:** 10.64898/2026.06.15.731303

**Authors:** Joe Wynn, Jochen Dierschke, Paul Dufour, Georg Langebrake, Robert E. Rollins, Heiko Schmaljohann, Pablo Salmon, Martin Irestedt, Sven Künzel, Christen M. Bossu, Kristen Ruegg, Anna Schnelle, Tianhao Zhao, Yong Chee Keita Sin, Franz Bairlein, Lars Burnus, Peter Hosner, Wieland Heim, Adriaan de Jong, Thiemo Karwinkel, Veronika Laine, Andreas Michalik, Micha Arved Neumann, Martin Päckert, Swen C. Renner, Markus Ünsold, Kevin Winker, Miriam Liedvogel

## Abstract

Vagrant animals – individuals found far outside their normal range – offer powerful natural experiments for understanding migratory mechanisms. The yellow-browed warbler (*Phylloscopus inornatus*) provides perhaps the best-yet example, typically migrating from Siberia to South/Southeast Asia yet found in increasingly large numbers in Western Europe. This represents a strikingly unresolved evolutionary puzzle: why do so many migrants consistently move in almost the complete wrong direction? A critical first step toward solving this enigma is determining where these birds come from. If vagrants came from the proximal western range edge this would imply simple disorientation, whilst a more easterly origin could imply large-scale ‘reverse’ misorientation. Here, we develop a geolocation-by-genotype algorithm for low-coverage whole-genome resequencing data collected from feathers. Our method identifies spatially informative SNPs; clusters them to account for covariance in allele frequency through space; and employs a bootstrapped maximum-likelihood framework to estimate spatial origin with uncertainty. Applied to more than 80 European-caught birds, our results place their origin in central Siberia (118°E; 89–134°E [95% CI]); over 2000km east of the western range edge. These results suggest mass misorientation in a near-reverse direction, and highlight the yellow-browed warbler as an exceptional system for probing the mechanism, ontogeny and evolution of migration.

## Introduction

Animal migration is one of the most remarkable adaptations to seasonal changes in the environment, allowing billions of animals to exploit seasonally occurring optimal resources and to avoid sub-optimal conditions by effecting coordinated long-distance movements (1). Such movements require the evolutionary fine-tuning of migratory propensity, timing and destination such that animals arrive in the correct place at the correct time, which in turn requires mechanisms capable of orchestrating and regulating predictable annual movements. Understanding such mechanisms is, therefore, not just a question of fundamental interest, but will be integral for understanding the extent to which migratory taxa will (or won’t) respond to rapidly occurring anthropogenic environmental alteration (2,3). Yet, studying the mechanisms of long-distance migration *in situ* is challenging, since experimental intervention often trades off against biological realism (4).

One solution is offered by ‘vagrant’ animals (individuals found outside their species’ normal range), which represent naturally occurring contrasts that can be used to tease apart migratory mechanisms without experimental manipulation. Vagrants likely arise through one or more of several processes: the very early stages of migratory route evolution; disorientation/misorientation caused by environmental factors; or errors in the innate programming or execution of long-distance migration (5–8). As such, vagrancy provides a powerful but underexploited paradigm for investigating the mechanism, ontogeny and evolution of animal migration (5).

Among vagrant systems, perhaps one of the most striking is the case of the yellow-browed warbler (*Phylloscopus inornatus*). This very small passerine (< 6.5 g) breeds across Siberia and typically winters in South and Southeast Asia, yet over the past half-century has become surprisingly and increasingly common in Western Europe, with thousands of birds now arriving each autumn at locations more than 5,000km from their ancestral non-breeding range (9). The scale of this phenomenon make it one of the most enigmatic evolutionary puzzles in the study of avian migration: how do such a large number of individuals of a single species come to migrate in the wrong direction, and does this represents disorientation, misorientation, or the early stages of a novel migratory route (9–11)?

A critical first step toward resolving this puzzle is to determine where yellow-browed warblers occurring in Europe originate within the vast Siberian breeding range. Under simple disorientation, where birds deviate randomly from their intended migratory direction, vagrants could in principle come from anywhere across the range, but individuals from the geographically proximal western edge would be expected to predominate since they face a shorter journey to Europe and hence a higher probability of survival (9). An origin concentrated further east, however, would be difficult to reconcile with random disorientation and would instead imply a large-scale misorientation (where birds move in a specific but inaccurate direction), demanding a fundamentally different mechanistic explanation (12). However, answering this question has proven exceptionally difficult. Laboratory-based orientation assays cannot identify the geographic origin of vagrant birds (10); Palearctic stable-isotope landscapes lack longitudinal resolution (13–16); extremely low adult survival of European-migrating individuals (< 1%) essentially prohibits the deployment of archival biologgers; and the species is in any case so small that contemporary tracking devices are of questionable utility when characterising natural behaviour.

Here, we overcome these limitations using a ‘genoscaping’ approach – mapping genetic variation through space; *sensu* Ruegg (17,18) – to determine the geographic origins of yellow-browed warblers caught in Europe. We develop a geolocation-by-genotype algorithm specifically designed for low-coverage whole-genome resequencing data mostly collected non-invasively from feathers, which incorporates a clustering and bootstrapping framework that accounts for correlations in allele frequencies between SNPs and allows the computation of meaningful confidence intervals. We apply this method to 82 European-caught individuals, including three historical museum specimens from 1873, and use the resulting geographic assignments to evaluate whether yellow-browed warbler vagrancy in Europe is best explained by disorientation from the range edge or by large-scale migratory misorientation.

## Methods

### Sample collection

European samples were collected on the Isle of Helgoland, Germany (n = 68; 7.88°E, 54.2°N); at Horumersiel, Germany (n = 6; 8.02°E, 53.7°N); on the Isle of Ouessant, France, (n = 5; - 5.07°E, 48.5°N); or at Edewecht, Germany (n = 1; 7.98°E, 53.1°N). The majority of samples were dropped feathers gathered during ringing activity, with exception of 3 toepad samples taken from birds collected in 1873 by the first warden of Helgoland, Heinrich Gätke, which were taken from specimens at the London Natural History Museum; and one tongue taken from a deceased bird on Ouessant. Siberian samples for georeferencing were collated from museum collections or via ringing activity from collections Europe (n = 31) or in the United States (n = 36). Samples were either tissue samples gathered during specimen collection (n = 37), toepads sampled from prepared specimens (n = 16), blood samples (n = 4), buccal swabs (n = 2) or feather samples taken from live birds during ringing activity (n = 91; see supplementary materials).

### DNA extraction, library preparation and sequencing

DNA was extracted from each tissue type using a protocol optimised for use with that particular tissue type. For pectoral muscle, blood and heart samples a Qiagen Puregene Precipitation-Based DNA Purification kit was used; for feather samples a salt extraction protocol using sodium acetate was used (19); and for toepads a Qiagen blood and tissue kit optimised for historical/ancient DNA was used (20).

Owing to the different properties of the different tissue types used in the analysis, libraries were prepared according to the different tissue types considered. Library preparation of historical DNA (hDNA) followed the procedures described by Irestedt et al. (21). In brief, libraries were prepared using the protocol of Meyer and Kircher (22), with modifications optimized for museum-derived samples (Irestedt et al., 2022). During the initial library preparation step, samples were treated with USER enzyme (New England Biolabs) to remove uracil residues and thereby reduce the characteristic deamination patterns associated with hDNA (23). After indexing (four dual-indexed libraries per individual), libraries were pooled and sequenced on the NovaSeq X Plus platform using one lane of the 10B-300 flow cell. Feather and tissue libraries were created using an adaptation of the protocol detailed in (24). Sequencing took place on two S4 lanes of an Illumina NovaSeq 3000 sequencing platform samples (n lane #1 = 62; n lane #2 = 73). 45 samples of other taxa were sequenced on lane #1 but are not presented here.

### Bioinformatics

Binary Alignment Map (BAM) files where created using a snakemake (25) pipeline for variant calling (26). In this pipeline adapters are trimmed using bbduk (27), duplicate sequences are removed using Samtools (28) and trimmed reads are mapped to the B10k yellow-browed warbler reference genome using bwa-mem2 (29).

To identify variants of interest from low-coverage WGS data, we used stringent filtering in ANGSD (30) based on recommendations in DeSaix et al. (17) and Lou et al. (31). Reads with a mapping quality > 30 and base quality > 33 were retained, and SNPs were retained based on a p-value of <1e^−6^. SNPs that had read data in at least 50% of individuals (n = 75), a minor allele frequency >0.05 and minimum and maximum total depths of 142 and 571 were retained. The minimum sequencing depth was determined as the minimum number of individuals required to call a variant (n = 75) multiplied by the mean sequencing depth of all individuals (75 individuals * 1.9x = 142); the maximum total depth was determined by 2 * total number of individuals * mean sequencing depth (2 * 150 * 1.9x = 571). The resultant genotype likelihoods were retained from ANGSD and used in downstream analysis.

### Geolocation using allele frequency

Previous attempts to geolocate samples by cross-referencing an individual’s genotype with distributions of allele frequency have sought to do so by summing likelihood estimates across all considered SNPs (32,33). Whilst the maximum likelihood estimates from such methods are completely interpretable, such approaches do not account for correlations in allele frequencies between SNPs – an inevitability, due to factors including linkage disequilibrium and the stochastic processes that lead to biogeographic patterns of allele frequencies (e.g. bottlenecks and founder effects) – which prohibits the calculation of accurate CIs during geographic assignment. This is because SNP-specific likelihoods would be assumed to be independent, which they are not, leading to massive overestimation of confidence. When dealing with low coverage sequencing data from a relatively sparse geographic sampling regime (see Figure 1c), quantifying model uncertainty is at least as important as accurate geolocation (if not more so). As such, we have updated existing methods to a) function using low-coverage data, and b) quantify uncertainty in estimates of location.

**Figure 1:**
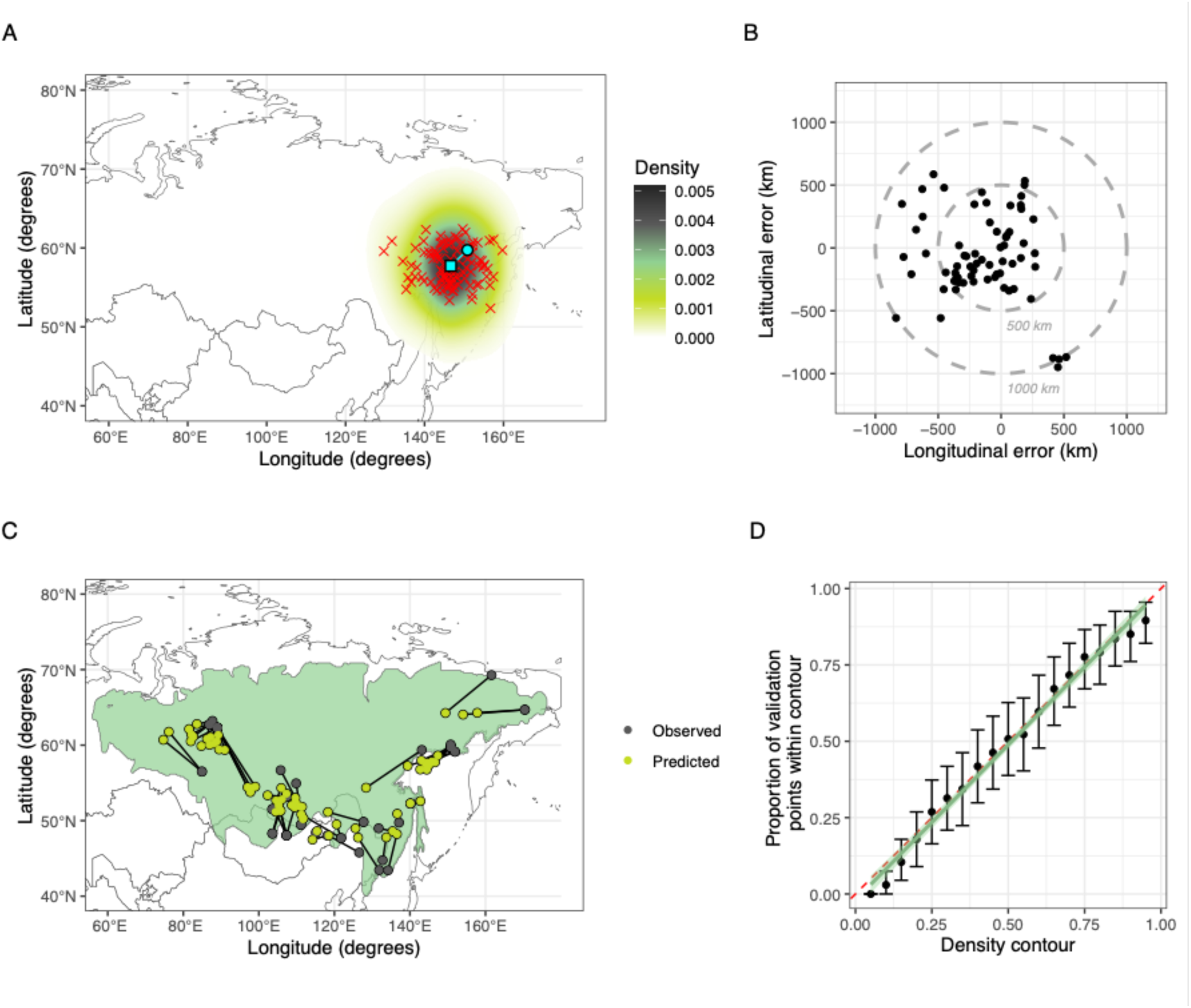
Validating the geolocation-by-genotype algorithm. A) An example geolocation of one validation datapoint. The true point is shown as a turquoise circle, the median estimated location as a turquoise square, with each iteration of the geolocation algorithm shown as a red cross. The background shows the 2D probability density. B) The longitudinal and latitudinal error in kilometres for each leave-one-out cross-validation (LOOCV) point. C) The estimated locations of points during LOOCV (predicted; green circles) linked to true locations (observed breeding locations; grey circles). Breeding range of the yellow-browed warbler is shown in green in the background. D) The proportion of LOO points that fall within a specific density contour value plotted against that contour value as a means to assessing how accurately our model quantified model uncertainty. The expected 1:1 line with a 0 intercept is shown as a red dashed line; the least mean squares regression line discussed in-text is shown as a green line (with green shading representing the standard error); with the bootstrapped 95% binomial confidence intervals per contour value shown as error bars.

All geolocation statistics were computed using R 4.5.1, and complete analysis scripts can be found in the supplementary materials of this submission. All of the below steps are illustrated in Figure S1.

### 1. Selecting informative SNPs and parameterising models of allele frequency

Not all SNPs will vary predictably through space, and not all variations through space can be picked up with the sparse sampling regime employed here. As such, we sought to isolate SNPs that varied detectably through space in a way that could be modelled accurately. These SNPs were isolated using four criteria, using a training dataset (80% of sequence data from known breeding sites in Siberia) to train models and a test dataset (20% of sequence data from known breeding sites in Siberia) to assess how well models fitted data they weren’t trained on.

#### Selection criteria #1

First, we sought to select SNPs that varied measurably through space by constructing quasibinomial generalised additive models (GAMs) of minor allele frequency through space. These models included linear predictive terms for longitude and latitude, as well as a ‘surface of a sphere’ spline with a basis dimension of five and a smoothing parameter of 0.5. This allowed allele frequencies to vary non-linearly, but meant that model complexity was fairly tightly constrained. This was seen as essential, since our sampling regime was relatively sparse (see Figure S2).

The total estimated number of copies of the minor allele at locus *i* for individual *j* (the ‘dosage’; here, *X*_*i*_*_j_*) was calculated as the as genotype likelihood L_ij_(g) multiplied by the number of alleles represented by that genotype:

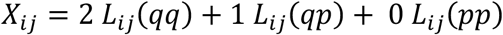

where *pp*, *pq*, and *qq* denote homozygous major, heterozygous, and homozygous minor genotypes, respectively. This dosage was divided by two to give the within-individual probability of any given allele being minor, which in turn gave a GAM with the following structure:

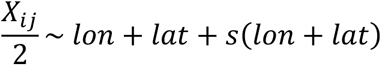

Where *s*(*lon* + *lat*) represents the spline of longitude and latitude described above. GAMs were fitted using the ‘mgcv’ package (34) with a quasibinomial error distribution and logit link to model 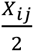. The ‘weights’ parameter was set to ‘2’ to indicate that 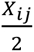 represented two Bernoulli trials per observation; and the quasibinomial family accounts for extra-binomial dispersion whilst allowing for non-integer *X*_*ij*_ values. SNPs were filtered to retain only those that performed significantly better than an intercept-only model in a likelihood ratio test (LRT; *⍺* = 0.05).

#### Selection criteria #2

Second, to ensure that variation in tissue type and/or sequencing coverage didn’t influence our results, we built a second model including these two potentially confounding variables alongside the spatial terms described above.

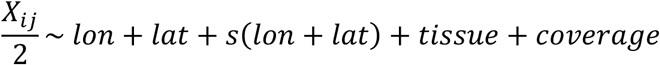

Tissue type was included as a potential confound since we had both contemporary fresh samples and museum toepad samples, with a much higher chance of de-amination and fragmentation expected in museum toepad samples owing to their age (35,36). Coverage was included to exclude sites where genotype uncertainty might influence the called genotype, and was calculated using the ‘CollectWgsMetrics’ function in GATK/Picard (37). SNPs were removed in instances where the model containing confounds fitted significantly better than the model containing only spatial terms (again, using an LRT; *⍺* = 0.05).

#### Selection criteria #3

The third filtering step involved testing the predictive ability of each model at each locus. For locus *i*, the GAM provided a model-predicted allele frequency *p*_*i*_ for the minor allele at a given longitude/latitude pair. Assuming Hardy–Weinberg equilibrium (HWE), the expected genotype frequencies 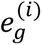 at locus *i* were then calculated as:

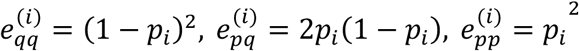

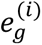 could then be compared to observed values outputted from ANGSD. For locus *i* and individual *j*, ANGSD provides genotype likelihoods L_ij_(g) = *P*(*D_*ij*_* | *g*), representing the probability of the observed sequencing data *D*_*ij*_ given each possible genotype *g* ∈ {*pp*, *pq*, *qq*}. This can be compared to 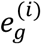 to assess model accuracy using the test portion of our dataset (i.e. the portion with which the model wasn’t trained). To do this we computed a Brier score (BS^(*i*)^) comparing the 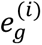 to L_ij_(g), calculating the BS^(*i*)^ per genotype *g* per individual *i* and then calculating a mean across genotypes and then *n* individuals.

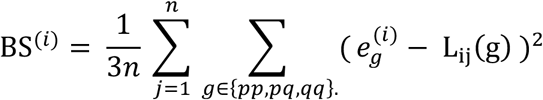

This allowed us to assess the goodness-of-fit of each allele frequency model, and we retained for analysis only alleles with a BS^(*i*)^< 0.05 (i.e. instances where the model predicted the test data with high accuracy).

#### Selection criteria #4

We also calculated the Brier Skill Score (*BSS*^(*i*)^) to eliminate models where high Brier scores were caused by extremely similar model predictions across all locations. To do this we calculated a baseline (reference) Brier score 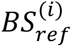 by randomising *p*_*i*_ between individuals before calculating 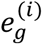 (and then the 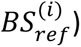), so as to create a baseline expectation of what the *BS*^(*i*)^ would be under an expectation of random predictions. We then calculated the *BSS*^(*i*)^ as:

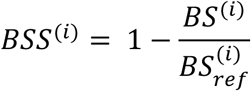

With *BSS*^(*i*)^ ≈ 1 meaning a far better fit than would be expected by chance; *BSS*^(*i*)^ ≈ 0 being no better than chance and *BSS*^(*i*)^ < 0 being that model fit was actively worse than chance. We then retained alleles with a *BSS*^(*i*)^ > 0.3 indicating a substantial improvement in model fit versus chance.

### 2. Allele spatial clustering

Based on the above subsetting, we reasoned that we retained only alleles that a) varied meaningfully through space; b) did not confound with tissue type; and c) predicted allele frequency (and hence genotype) with accuracy beyond that expected y chance. This allowed us to filter 10,970,393 SNPs (retained by our ANGSD variant calling pipeline) down to 16,238 SNPs that varied through space in a manner that we could accurately model.

For each of these SNPs we used the GAMs described above to estimate allele frequency for a 5° x 5° grid extending from 60°E, 40°N in the south-west to 180°E, 70°N in the north-east. We then performed principal component analysis (PCA) to assess how similar different SNPs were in the way their minor allele frequency varied through space. We did this by assigning each allele to one of *k* clusters using k-means clustering based on the first three principal components (PCs), with these PCs chosen as they cumulatively explained >95% of total variance. We tested several different *k* values ranging from 10 to 200, and used an elbow plot of the within-cluster sum-of-squares to find the point at which adding more clusters did not meaningfully increase model fit. We found this point to be 150 clusters (see Figure S3). We did not, however, assume that this cluster value was accurate, since even slight deviations from the true value would affect whether or not we could calculate informative confidence intervals. Instead, we directly tested whether confidence intervals were computed appropriately in our validation dataset (see below).

### 3a. Geolocation via maximum likelihood: combining likelihoods from different SNPs

Since we found that alleles were distributed in 150 clusters, we reasoned that calculating joint likelihood from all 16,238 SNPs simultaneously would lead to massive overestimation of likelihood and hence – should we want to calculate confidence intervals – these would likely be hugely underestimated. As such, we randomly sampled one locus per cluster per iteration of our geolocation algorithm. We then calculated a maximum likelihood value based on the 150 SNPs considered using the below method, before repeating the process with a new set of SNPs (again, with one locus sampled per cluster). We then stored the estimated locations per iteration, calculating the most likely location as the median bootstrapped longitude/latitude and 95% confidence intervals as the 0.025 and 0.975 contours in a 2D density kernel (bandwidth = 5; see Figure 1). Importantly, for each focal individual we only considered SNPs for which the focal individual had sequencing data (i.e. the ANGSD genotype was not 0.333/0.333/0.333). This ensured that only informative SNPs were considered per iteration.

### 3b. Geolocation via maximum likelihood: geolocation optimisation algorithm

The most likely location for each individual per iteration was calculated using a ‘Limited-memory Broyden–Fletcher–Goldfarb–Shanno algorithm with Bounds’ (L-BFGS-B) implemented using the ‘optim’ function in R, fitted with bounds of −180° and 180° for longitude and −90° and 90° for latitude.

As noted above, for locus *i* and individual *j* ANGSD provides genotype likelihoods L_ij_(g) = *P*(*D*_*ij*_ ∣ *g*). Since our model specifies the expected probabilities of these genotypes at locus *i*, denoted 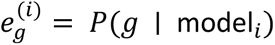, we were able to geolocate samples by comparing ANGSD output to our model predictions. Because the true genotype is unknown, we marginalized over all possible genotypes to obtain the probability of the observed data given the model:

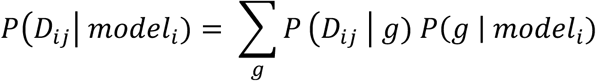

This expression integrates over the latent genotype state, weighting the likelihood of the observed data under each genotype by the model’s predicted genotype frequency. Assuming independence between SNPs, which we reasoned to be valid given our above clustering step, the overall likelihood of the observed dataset under the model is the product of these per-observation probabilities:

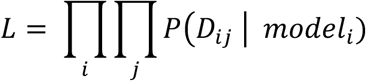

Or, for numerical stability and to prevent underflow:

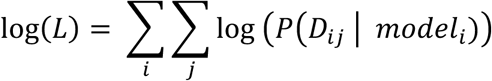

The resulting log-likelihood provides a measure of model fit that accounts for uncertainty in individual genotype calls, which in turn can be used in an optimisation algorithm to find the most likely location based on genotype data at each locus for a given individual. The negative of this was optimised using L-BFGS-B, with the resultant estimated longitude/latitude used in our model averaging/pseudo-bootstrapping calculations (see above).

### Assessing geolocation-by-genotype model efficacy

Assessing the efficacy of our geolocation-by-genotype algorithm is essential before applying it to new data, especially since we were using low coverage sequencing data gathered with a comparatively sparse sampling regime from across the breeding range. We wanted to assess, therefore, whether a) our model could geolocate accurately and b) whether our model could parameterise uncertainty in its estimates accurately.

To do this we used leave-one-out cross-validation (LOOCV), a form of *k*-fold cross validation where the model is trained on all-but-one samples, with this sample kept back to test model efficacy. We used LOOCV since this allowed for the model to be as representative as possible of how it would function on ‘real’ data, whilst not allowing for over-fitting by including test datapoints in the training data. Following LOOCV for each of our georeferenced samples we used the ‘distHaversine’ function from the ‘geosphere’ package in R (38) to calculate the distance from the estimated to the true location. These we used to assess model accuracy, calculating median, mean, standard deviation and maximum error. We also performed a randomisation analysis, repeatedly randomising predicted location with respect to true location and calculating the mean error under an assumption of random prediction. We then compared the true error to this distribution in a two-tailed permutation analysis to assess whether our model performed better than would be expected by chance.

To assess how well our model quantified uncertainty, we assessed whether the true locations extracted from LOOCV fell inside the *n^th^* quantile, with the expectation that the overall proportion of points that fell within the *n^th^* density contour should be *n*. For example, we would expect the true point to fall within the 95% density contour 95% of the time; the true point to tall within the 50% density contour 50% of the time; and so on. Quantiles were constructed using the ‘kde2d’ function from the ‘MASS’ package in R (39), with the *n^th^* density contour identified using the ‘contourLines’ function. We constructed a linear regression between the quantile values (varying from 5% to 95% in 5% increments) and the proportion of times the LOO validation points fell within that contour, with the *a priori* expectation that – should our model quantify uncertainty correctly – this should produce a line with a gradient of 1.0; an intercept of 0.0; and an extremely high r-squared value.

### Using our geolocation-by-genotype model

Our model was run on European-caught samples for 100 iterations, each time resampling one allele per cluster as a quasi-bootstrap (see above), from which the most likely longitude/latitude ± CIs were calculated per bird (see Figure 2a). The most likely mean location was calculated as the cell with the highest density in the sum of all 2D density kernels constructed per bird; with 95% CIs calculated as the minimum/maximum longitude/latitude values reported in the 95% contour of this density kernel (see Figure 2b).

**Figure 2:**
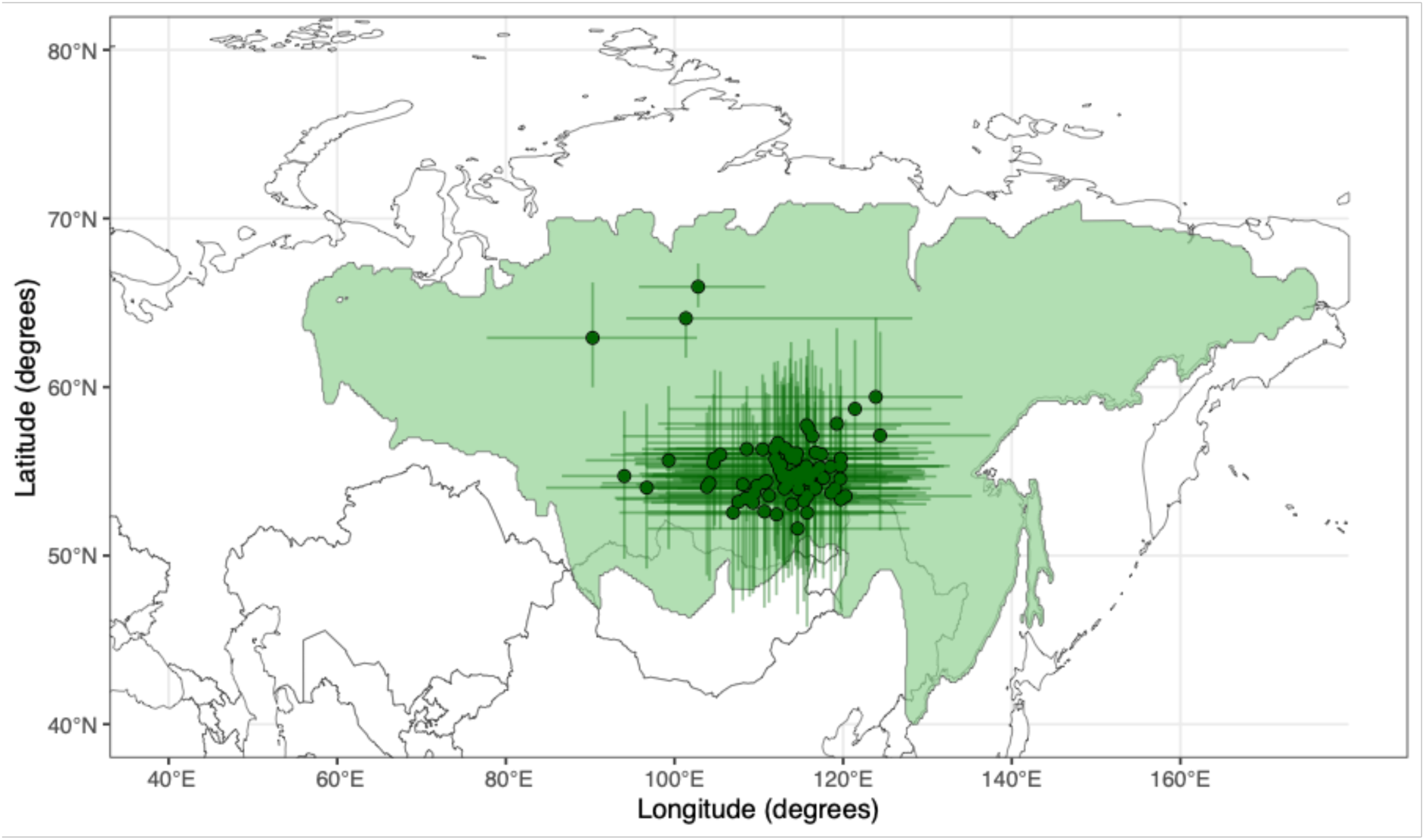
The origin of European-caught yellow-browed warblers. Median estimated locations (green points) ± 95% CI (green error bars) for yellow-browed warblers caught during winter in Europe, with background shading denoting the yellow-browed warbler breeding range.

## Results

To ensure our model behaved as expected, we used leave-one-out cross-validation (LOOCV) on WGS data from birds where the sampling site was known to assess model performance. We found that assignment functioned with a mean error of 426 km (± 261km [SD]), a median error of 352 km and a maximum error of 1077 km, with the model performing far better than would be expected by random chance (randomisation; p < 0.0001; see Figure 1b and Figure 1c). Given that the yellow-browed warbler breeding range spans > 5000 km, this suggests that our algorithm was capable of estimating geographic origin with reasonable accuracy. We also estimated the number of times that the true position fell within different 2D density contours (5% - 95%, at 5% intervals), with the expectation that the proportion of instances where the true point fell within the contour in question should be equal to the contour value (e.g. the true location should fall within the 95% contour 95% of the time). We found a near-perfect correlation (r^2^ = 0.990) between the expected and observed proportions, with a gradient that was almost exactly 1 and an intercept of almost exactly 0 (gradient = 1.01 ± 0.0257 [SE]; intercept = 0.019 ± 0.0147 [SE]; ANOVA F = 1,559, p < 0.001; see Figure 1d). This – when taken alongside estimates of accuracy – suggests that our algorithm is a) capable of geographic assignment with reasonable accuracy, and b) capable of quantifying uncertainty appropriately.

Having established the efficacy of our algorithm on georeferenced breeding samples, we next applied it to the low-coverage WGS data from yellow-browed warblers caught across Western Europe. Surprisingly, we found that all European-caught yellow-browed warblers – including both modern (2011-2021; n = 79) and ancient samples (1873; n = 3) – did not come from the western range edge. Instead, we found that birds from both Germany and France originated from a fairly constrained location – a maximum likelihood position of 118.2°E (88.9°E – 133.9°E [95% CI]), 55.2°N (46.1°N – 67.2°N [95% CI]; see Figures 2 and 3) – east of Lake Baikal in central Siberia. Whilst geographic assignment with low-coverage genomic data is necessarily a noisy process, our results seemingly exclude the possibility that birds come from the western edge of the range in the Ural Mountains (something that could not be ruled out using isotopic analysis; (13)).

**Figure 3:**
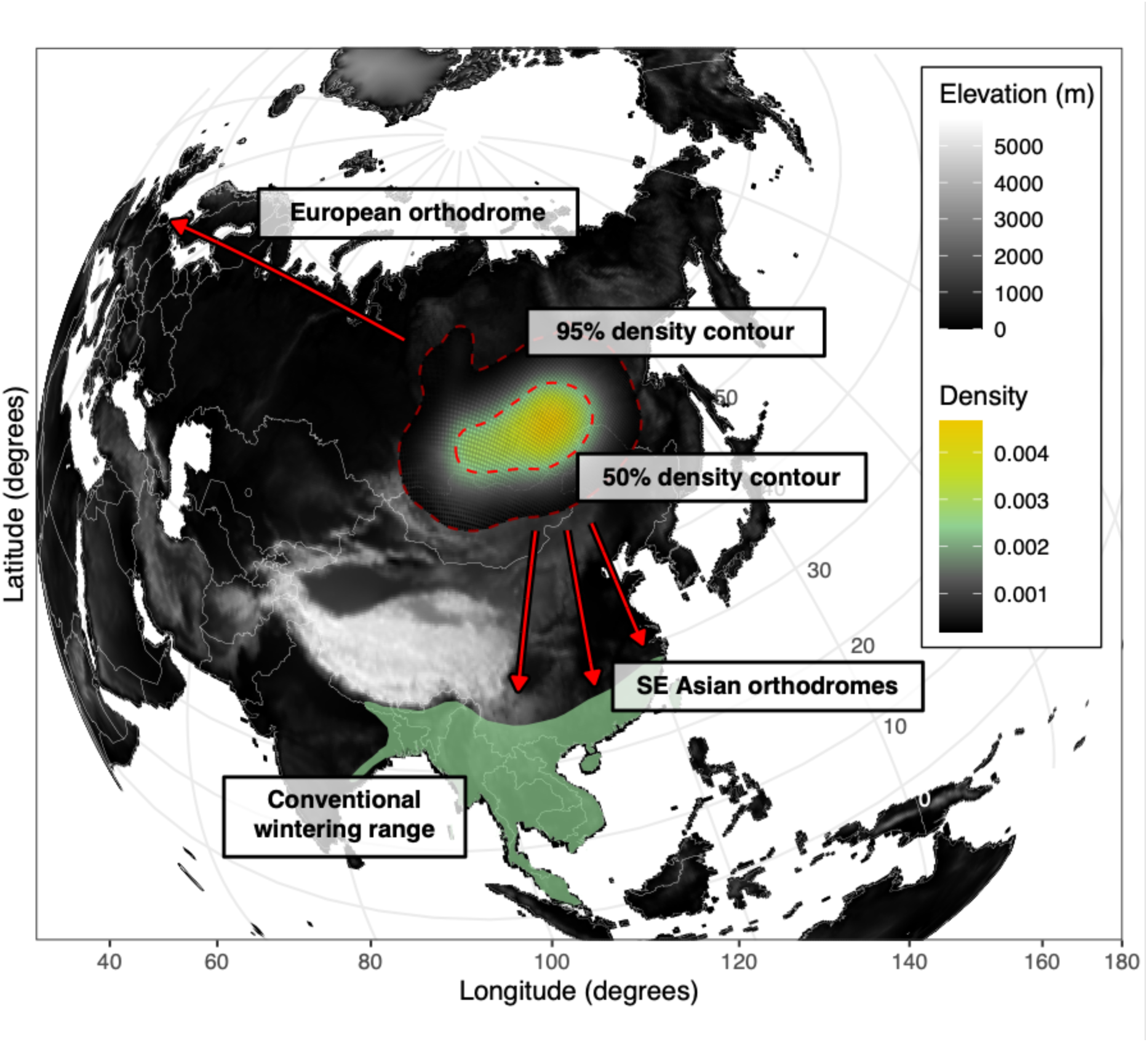
The geometry of yellow-browed warbler reorientation. The likely origins of European-caught yellow-browed warblers are shown as a 2D density plot, with 95% (dark red) and 50% (light red) density contours denoted with text. The conventional wintering range in South and Southeast Asia is highlighted in green; the background shading denotes altitude; and loxodromes (i.e. Great Circle routes) from the most likely mean origin location are shown extending to both Helgoland and to multiple parts of the conventional wintering range. Owing to the spherical projection used, coordinate system doesn’t intersect the y-axis. As such, latitudinal degrees are shown on the right of the plot along a gridline.

## Discussion

Using whole-genome resequencing data, we develop a model capable of estimating the breeding locations of vagrant birds caught in Europe. This model builds on existing solutions to function using low-coverage data and a comparatively sparse sampling regime, whilst also incorporating the calculation of confidence intervals. Using this model, we geolocate European-caught yellow-browed warblers to breeding sites in central Siberia; far further east than previously expected (13). As with any geolocation technology, it is possible that the location estimate derived is biased by some as-yet-unknown process. However, given that the model in question function as expected during *k*-fold cross-validation, we suggest that the most parsimonious explanation of our results is that European-caught yellow-browed warbler do not come from their range edge.

Since European-migrating yellow-browed warblers do not come from the geographically proximal western part of the species’ range, we propose that – consistent with long-held speculation (9–11) – it is unlikely that the mechanism underlying yellow-browed warbler vagrancy in Europe is simple disorientation (i.e. random movements owing to a lack of positional/directional information). This is because such processes would necessarily favour those starting the shortest distance from Western Europe. Instead, we suggest that an as-yet-unknown process drives a reorientation in almost completely the wrong direction (see Figure 3) for birds in a surprisingly constrained subset of the yellow-browed warbler breeding population (10). In developmental biology, key discoveries relating to inheritance, ontogeny and mechanism are often discovered owing to pronounced changes in phenotype in well characterised model systems (e.g. *eyeless*; (40,41). In much the same way, we posit that the extreme reorientation observed in yellow-browed warblers provides an analogous opportunity to probe the mechanism and ontogeny of long-distance avian migration. What might such a drastic change in orientation represent?

### Day-to-day navigational errors caused by topography and the environment

Topographic barriers (e.g. the Qinghai-Tibet Plateau) and/or environmental/physical conditions (e.g. magnetic anomalies) en route could cause birds to reorient in a very specific way (42). This might seem the least likely explanation, since it is unclear how such errors produce targeted misorientation rather than random disorientation. Nonetheless, given how imperfect our understanding of the mechanisms of long-distance migration is (4,43), the possibility of topography/physical distortions causing severe misorientation should not be discounted out-of-hand.

**Ontogenetic conditions:** it is thought that a period of early life, phase-specific learning allows birds to a) learn the spatial cues associated with the breeding site (44,45) and b) calibrate the various compass cues that will eventually guide long-distance migration (46,47). It is possible that some part of these processes functions imperfectly in Siberia – e.g. owing to proximity to the zero isoline of magnetic declination line (located at around 98°E at this latitude; (48)) – giving rise to surprising and highly divergent variation in orientation preference.

**Differences in genomic information:** it is well known that night-migratory songbirds rely on genetic information to guide outbound migration, though how this information is genetically encoded is not fully understood (49–52). It is possible that *de novo* point mutations, structural variation (e.g. LTR insertions) or epistatic interactions acting on standing variation could all produce aberrant phenotypes, all of which could drive migration of yellow-browed warblers to Europe. This has long been suggested to be the case in the Eurasian blackcap (*Sylvia atricapilla*), where birds have been shown to increasingly favour a heritable ‘inverted’ northwards migratory route over their traditional southern migration (53–56). This is strikingly similar to the changes in migration observed in yellow-browed warblers, hence it is tempting to suggest that a genetic explanation might be invoked here also.

Given the above, it might seem possible that an increasing number of yellow-browed warblers in Europe represents the early stages of migratory route evolution. However, unlike in other Siberian species that occur in Europe (6,8), between-year yellow-browed warbler ringing recoveries are extremely rare (57) and the number of adult yellow-browed warblers observed vanishingly small (<c.1-5%; (16). This is in contrast even with the closely-related Siberian chiffchaff (*Phylloscopus collybita tristis;* another Siberian vagrant found in Europe), where the proportion of adult birds seems almost an order of magnitude higher (16). Hence, even accounting for the r-selected life history of *Phylloscopus* warblers, this might make evolution the less likely explanation of increased numbers of vagrant yellow-browed warblers.

If this were true, the recent increase in the number of yellow-browed warblers would not be explained by selection in favour of reorienting birds, but would instead be caused by other factors. For instance: changing climatic conditions encountered en route increasing survival rates; or an increased population size at the breeding range (since a fixed proportion of reorienting birds will produce more vagrants from a larger population). Indeed, yellow-browed warblers reach highest densities in recently burned taiga forest patches (58), which are more common in central Siberia in recent years owing to changes in the climate (59). Therefore, it is not necessarily a given that year-on-year variation in the number of European yellow-browed warblers is linked directly to the cause of their reorientation.

Selecting between and favouring any explanations of reorientation based on the sparse available information is of questionable utility. Nonetheless, we believe that our findings represent the first evidence that vagrant yellow-browed warblers are not simply overspilling individuals from the range edge, and instead represent migratory reorientation on a truly impressive scale. We also propose that a better understanding of the genomic basis of migration will help us to elucidate this enigmatic system still further, and that in turn the mass reorientation of yellow-browed warblers will be of considerable utility when considering our understanding of the mechanism, development and inheritance of avian migratory phenotypes. In a more general sense, we also suggest these methods, and particularly our updated geolocation-by-genotype algorithm for use with low-coverage sequencing data, might represent a powerful tool in the study of ecology more generally – particularly in the study of movement and dispersal of a wide variety of taxa over a large range of spatial scales.

## Acknowledgments

We thank all the ornithologists who contributed to the infrastructure that enabled the sampling regime used in this project.

## Author contributions

Conceptualization: JW, JD, ML

Data curation: JW, PD, HS, FB, LB, PH, WH, AdJ, VL, AM, MAN, MP, SCR, MÜ, KW

Methodology: JW, ML, PD, GL, RER, PS, MI, SV, CMB, KR, AS, TZ

Investigation: JW, ML, PD, GL

Visualization: JW

Funding acquisition: JW, ML

Project administration: JW, PD, ML

Supervision: JW, ML

Writing – original draft: JW

Writing – review & editing: JW, JD, YCKS, PD, GL, RER, PS, HS, MI, SV, CMB, KR, AS, TZ, FB, LB, PH, WH, AdJ, VL, AM, MAN, MP, SCR, MÜ, KW, ML

## Data, code, and materials availability

All data, code, and materials used in the analysis are available at:

https://zenodo.org/records/18518858?preview=1&token=eyJhbGciOiJIUzUxMiJ9.eyJpZCI6ImQ1OGFjMjg1LWE2YmUtNDM5NS1hODJhLWQyMzEwNWVlNWRhNCIsImRhdGEiOnt9LCJyYW5kb20iOiJlYTE1MDZlZTZiODVlMzk3MDk5MGEwMTYyMmU5YTg4YyJ9.GZyQFsoNb13gFm6DB0v4LPbgRuQxg90VCqdkvvrHw8UFvEVSLjZb-qlkaJTl4OrViHSl6dGoMWtqyST8V4fI6w

---

**Figure S1:**
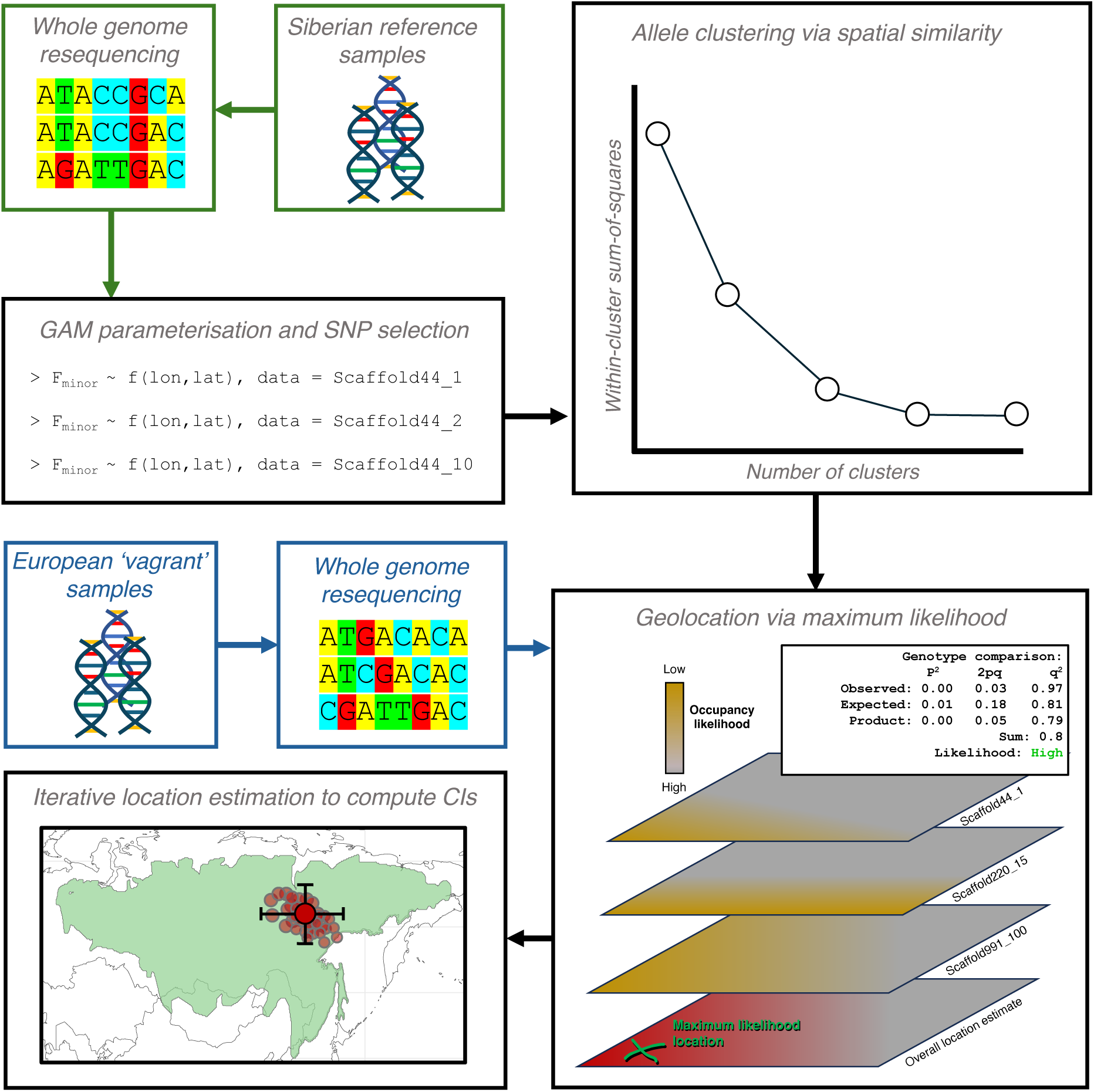
The geolocation-by-genotype method. A flow chart showing how georeferenced samples (highlighted in green) are used to model allele frequency through space, and how these models are then integrated with vagrant samples (highlighted in blue) are to calculate the range of possible locations the vagrant samples might come from.

**Figure S2:**
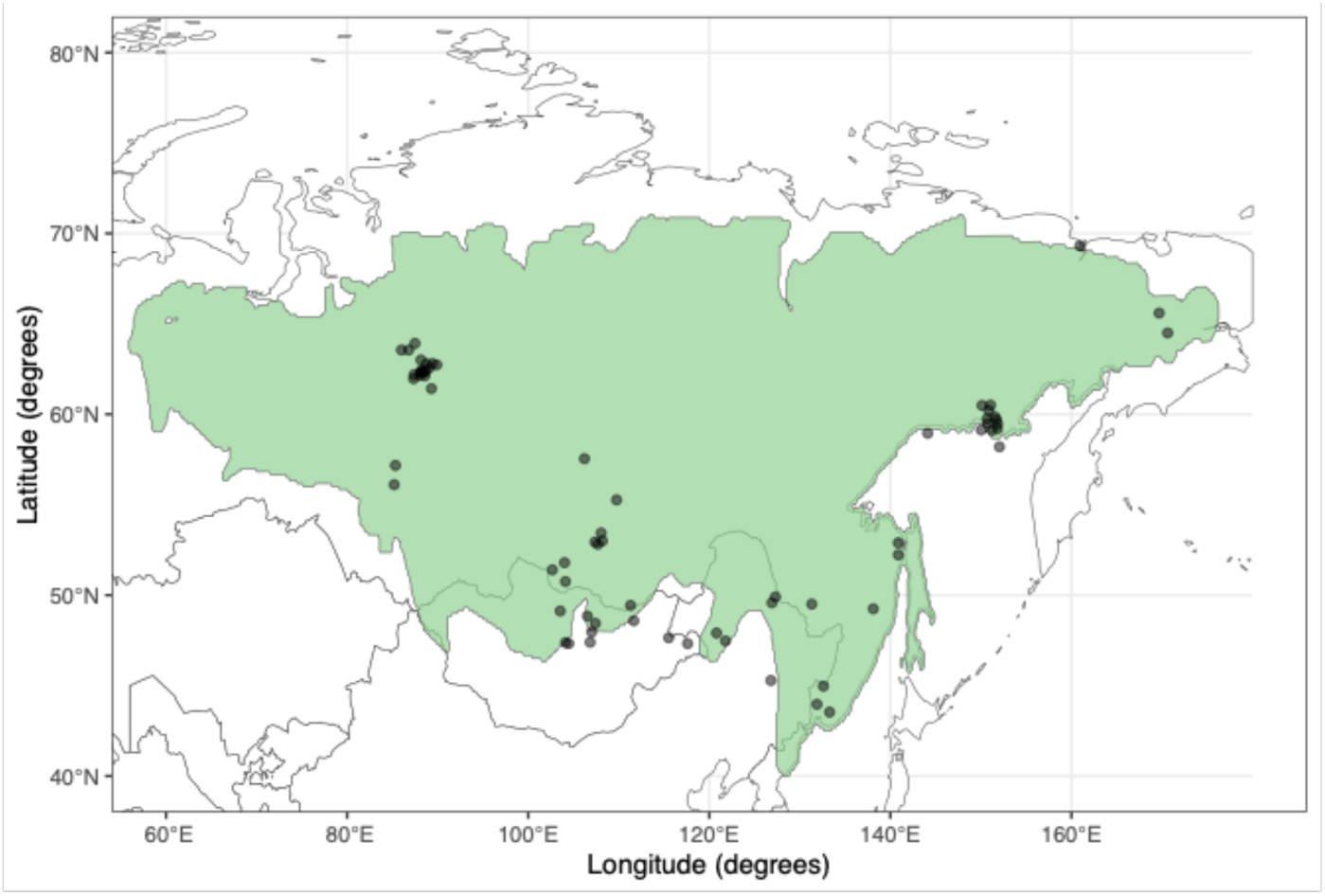
Sampling regime of georeferenced breeding yellow-browed warbler used in our analysis. The breeding range of the yellow-browed warbler is shown in light green, and a small amount of jitter (0.1°) has been induced in the x/y coordinates of each point to aid with visualising the number of points sampled at each location.

**Figure S3:**
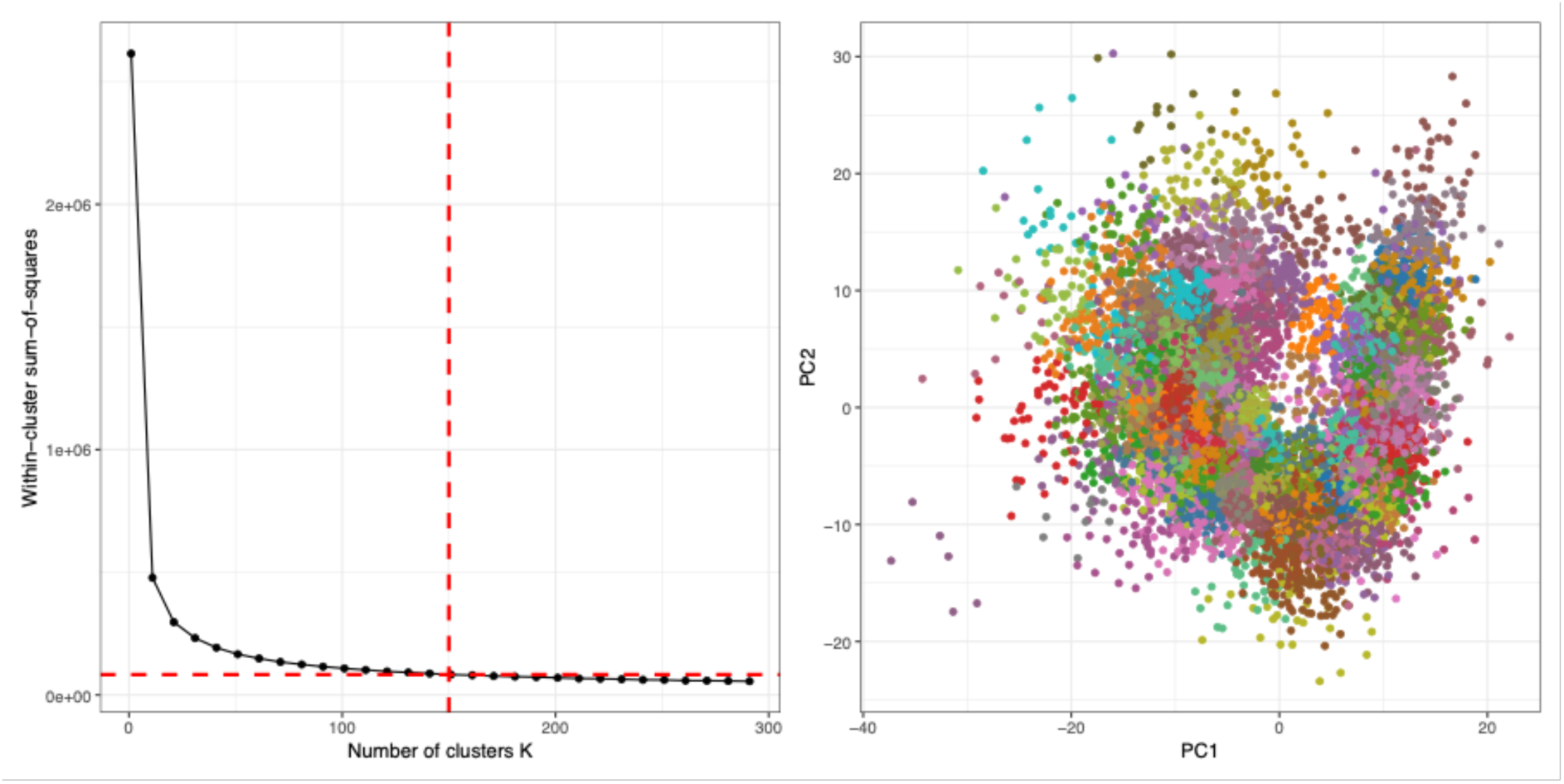
Iterative resampling as a solution to achieve non-independence in location estimates. A) Elbow plot of within-cluster sum-of-squares for 0 to 200 SNP clusters, indicating that 150 clusters represents the point at which adding further clusters does not meaningfully increase variance explained. B) PC1 and PC2 of allele frequency spatial variance coloured by cluster ID for each of our 150 clusters.

## References

1. Winger BM, Auteri GG, Pegan TM, Weeks BC. A long winter for the Red Queen: rethinking the evolution of seasonal migration. Biol Rev. 2019;94(3):737–52. doi:10.1111/brv.12476

2. Linssen H, van Loon EE, Shamoun-Baranes JZ, Nuijten RJM, Nolet BA. Migratory swans individually adjust their autumn migration and winter range to a warming climate. Glob Change Biol. 2023;29(24):6888–99. doi:10.1111/gcb.16953

3. Lewin PJ, Wynn J, Arcos JM, Austin RE, Blagrove J, Bond S, et al. Climate change drives migratory range shift via individual plasticity in shearwaters. Proc Natl Acad Sci U S A. 2024 Feb 6;121(6):e2312438121. doi:10.1073/pnas.2312438121 PubMed PMID: 38285933; PubMed Central PMCID: PMC10861922.

4. Wynn J, Liedvogel M. Lost: on what level should we aim to understand animal navigation? J Exp Biol. 2023 May 26;226(10):jeb245441. doi:10.1242/jeb.245441

5. Dufour P, Lees AC, Gilroy J, Crochet PA. The overlooked importance of vagrancy in ecology and evolution. Trends Ecol Evol. 2024 Jan 1;39(1):19–22. doi:10.1016/j.tree.2023.10.001 PubMed PMID: 37945456.

6. Dufour P, Franceschi C de, Doniol-Valcroze P, Jiguet F, Guéguen M, Renaud J, et al. A new westward migration route in an Asian passerine bird. Curr Biol. 2021 Dec 20;31(24):5590–5596.e4. doi:10.1016/j.cub.2021.09.086 PubMed PMID: 34687610.

7. Lees A, Gilroy J. Vagrancy in Birds. Helm; 2021.

8. Lees AC, Gilroy JJ. Bird migration: When vagrants become pioneers. Curr Biol. 2021 Dec 20;31(24):R1568–70. doi:10.1016/j.cub.2021.10.058 PubMed PMID: 34932963.

9. Dufour P, Åkesson S, Hellström M, Hewson C, Lagerveld S, Mitchell L, et al. The Yellow-browed Warbler (Phylloscopus inornatus) as a model to understand vagrancy and its potential for the evolution of new migration routes. Mov Ecol. 2022 Dec 14;10(1):59. doi:10.1186/s40462-022-00345-2

10. Thorup K. Vagrancy of Yellow-browed Warbler Phylloscopus inornatus and Pallas’s Warbler Ph. proregulus in north-west Europe: Misorientation on great circles? Ringing Migr. 1998 May 1;19(1):7–12. doi:10.1080/03078698.1998.9674155

11. Rabøl J. Reversed migration as the cause of westward vagrancy by four Phylloscopus warblers. Br Birds. 1969;62:89–92.

12. Thorup K. Reverse migration as a cause of vagrancy. Bird Study. 2004 Oct 1;51(3):228–38. doi:10.1080/00063650409461358

13. Jong A, Torniainen J, Bourski OV, Heim W, Edenius L. Tracing the origin of vagrant Siberian songbirds with stable isotopes: the case of Yellow-browed Warbler (Abrornis inornatus) in Fennoscandia. Ornis Fenn. 2019 Jul 1;96(2):90–9. doi:10.51812/of.133950

14. Rocque DA, Ben-David M, Barry RP, Winker K. Assigning birds to wintering and breeding grounds using stable isotopes: lessons from two feather generations among three intercontinental migrants. J Ornithol. 2006 Apr 1;147(2):395–404. doi:10.1007/s10336-006-0068-2

15. Dufour P, Kardynal KJ, Hobson KA, Monticelli D, Kolbeinsson Y, Alfrey P, et al. Origins of Nearctic migratory landbird vagrants recorded in Europe revealed by feather isotopic analysis. Sci Rep. 2025 May 2;15(1):15456. doi:10.1038/s41598-025-99765-4

16. Dufour P, Hellström M, Franceschi C de, Illa M, Norevik G, Cuchot P, et al. Using age-ratios to investigate the status of two Siberian Phylloscopus species in Europe. Ibis. 2025;167(3):632–45. doi:10.1111/ibi.13382

17. DeSaix MG, Anderson EC, Bossu CM, Rayne CE, Schweizer TM, Bayly NJ, et al. Low-coverage whole genome sequencing for highly accurate population assignment: Mapping migratory connectivity in the American Redstart (Setophaga ruticilla). Mol Ecol. 2023;32(20):5528–40. doi:10.1111/mec.17137

18. Ruegg KC, Harrigan RJ, Saracco JF, Smith TB, Taylor CM. A genoscape-network model for conservation prioritization in a migratory bird. Conserv Biol. 2020;34(6):1482–91. doi:10.1111/cobi.13536

19. Aljanabi SM, Martinez I. Universal and rapid salt-extraction of high quality genomic DNA for PCR-based techniques. Nucleic Acids Res. 1997 Nov 1;25(22):4692–3. doi:10.1093/nar/25.22.4692

20. Latz MAC, Grujcic V, Brugel S, Lycken J, John U, Karlson B, et al. Short- and long-read metabarcoding of the eukaryotic rRNA operon: Evaluation of primers and comparison to shotgun metagenomics sequencing. Mol Ecol Resour. 2022;22(6):2304–18. doi:10.1111/1755-0998.13623

21. Irestedt M, Thörn F, Müller IA, Jønsson KA, Ericson PGP, Blom MPK. A guide to avian museomics: Insights gained from resequencing hundreds of avian study skins. Mol Ecol Resour. 2022;22(7):2672–84. doi:10.1111/1755-0998.13660

22. Meyer M, Kircher M. Illumina Sequencing Library Preparation for Highly Multiplexed Target Capture and Sequencing. Cold Spring Harb Protoc. 2010 Jan 6;2010(6):pdb.prot5448. doi:10.1101/pdb.prot5448 PubMed PMID: 20516186.

23. Briggs AW, Stenzel U, Meyer M, Krause J, Kircher M, Pääbo S. Removal of deaminated cytosines and detection of in vivo methylation in ancient DNA. Nucleic Acids Res. 2010 Apr 1;38(6):e87. doi:10.1093/nar/gkp1163

24. Schweizer TM, DeSaix MG. Cost-effective library preparation for whole genome sequencing with feather DNA. Conserv Genet Resour. 2023 Jun 1;15(1):21–8. doi:10.1007/s12686-023-01299-2

25. Mölder F, Jablonski KP, Letcher B, Hall MB, Dyken PC van, Tomkins-Tinch CH, et al. Sustainable data analysis with Snakemake [Internet]. F1000Research; 2025 [cited 2025 Dec 4]. Available from: https://f1000research.com/articles/10-33 doi:10.12688/f1000research.29032.3

26. Langebrake G, matwe340. gmanthey/variant-calling: Release version 1.0.0 [Internet]. Zenodo; 2025 [cited 2025 Dec 4]. Available from: https://zenodo.org/records/15969023 doi:10.5281/zenodo.15969023

27. SourceForge [Internet]. 2025 [cited 2025 Dec 4]. BBMap. Available from: https://sourceforge.net/projects/bbmap/

28. Danecek P, Bonfield JK, Liddle J, Marshall J, Ohan V, Pollard MO, et al. Twelve years of SAMtools and BCFtools. GigaScience. 2021 Feb 1;10(2):giab008. doi:10.1093/gigascience/giab008

29. Vasimuddin Md, Misra S, Li H, Aluru S. Efficient Architecture-Aware Acceleration of BWA-MEM for Multicore Systems. In: 2019 IEEE International Parallel and Distributed Processing Symposium (IPDPS) [Internet]. 2019 [cited 2025 Dec 4]. p. 314–24. Available from: https://ieeexplore.ieee.org/document/8820962 doi:10.1109/IPDPS.2019.00041

30. Korneliussen TS, Albrechtsen A, Nielsen R. ANGSD: Analysis of Next Generation Sequencing Data. BMC Bioinformatics. 2014 Nov 25;15(1):356. doi:10.1186/s12859-014-0356-4

31. Lou RN, Therkildsen NO. Batch effects in population genomic studies with low-coverage whole genome sequencing data: Causes, detection and mitigation. Mol Ecol Resour. 2022;22(5):1678–92. doi:10.1111/1755-0998.13559

32. Rañola JM, Novembre J, Lange K. Fast spatial ancestry via flexible allele frequency surfaces. Bioinformatics. 2014 Oct 15;30(20):2915–22. doi:10.1093/bioinformatics/btu418

33. Guillot G, Jónsson H, Hinge A, Manchih N, Orlando L. Accurate continuous geographic assignment from low- to high-density SNP data. Bioinformatics. 2016 Apr 1;32(7):1106–8. doi:10.1093/bioinformatics/btv703

34. Wood S. mgcv: Mixed GAM Computation Vehicle with Automatic Smoothness Estimation [Internet]. 2000 [cited 2025 Nov 8]. p. 1.9–4. Available from: https://CRAN.R-project.org/package=mgcv doi:10.32614/CRAN.package.mgcv

35. Settlecowski AE, Marks BD, Manthey JD. Library preparation method and DNA source influence endogenous DNA recovery from 100-year-old avian museum specimens. Ecol Evol. 2023;13(8):e10407. doi:10.1002/ece3.10407

36. Sproul JS, Maddison DR. Sequencing historical specimens: successful preparation of small specimens with low amounts of degraded DNA. Mol Ecol Resour. 2017;17(6):1183–201. doi:10.1111/1755-0998.12660

37. McKenna A, Hanna M, Banks E, Sivachenko A, Cibulskis K, Kernytsky A, et al. The Genome Analysis Toolkit: A MapReduce framework for analyzing next-generation DNA sequencing data. Genome Res. 2010 Jan 9;20(9):1297–303. doi:10.1101/gr.107524.110 PubMed PMID: 20644199.

38. Hijmans RJ. geosphere: Spherical Trigonometry [Internet]. 2010 [cited 2025 Nov 8]. p. 1.5–20. Available from: https://CRAN.R-project.org/package=geosphere doi:10.32614/CRAN.package.geosphere

39. Ripley B, Venables B. MASS: Support Functions and Datasets for Venables and Ripley’s MASS [Internet]. 2009 [cited 2025 Nov 8]. p. 7.3–65. Available from: https://CRAN.R-project.org/package=MASS doi:10.32614/CRAN.package.MASS

40. Hoge MA. Another Gene in the Fourth Chromosome of Drosophila. Am Nat. 1915 Jan;49(577):47–9. doi:10.1086/279455

41. Quiring R, Walldorf U, Kloter U, Gehring WJ. Homology of the eyeless Gene of Drosophila to the Small eye Gene in Mice and Aniridia in Humans. Science. 1994 Aug 5;265(5173):785–9. doi:10.1126/science.7914031

42. Gulson-Castillo ER, Van Doren BM, Bui MX, Horton KG, Li J, Moldwin MB, et al. Space weather disrupts nocturnal bird migration. Proc Natl Acad Sci. 2023 Oct 17;120(42):e2306317120. doi:10.1073/pnas.2306317120

43. Guilford T, Åkesson S, Gagliardo A, Holland RA, Mouritsen H, Muheim R, et al. Migratory navigation in birds: new opportunities in an era of fast-developing tracking technology. J Exp Biol. 2011 Nov 15;214(22):3705–12. doi:10.1242/jeb.051292

44. Wynn J, Padget O, Mouritsen H, Morford J, Jaggers P, Guilford T. Magnetic stop signs signal a European songbird’s arrival at the breeding site after migration. Science. 2022 Jan 28;375(6579):446–9. doi:10.1126/science.abj4210

45. Wynn J, Oliver Padget, Mouritsen H, Perrins CM, Guilford TC. Natal imprinting to the Earth’s magnetic field in a pelagic seabird: Current Biology. Curr Biol. 2020;30(14):2869–73.

46. Alerstam T, Högstedt G. The role of the geomagnetic field in the development of birds’ compass sense. Nature. 1983 Dec;306(5942):463–5. doi:10.1038/306463a0

47. Emlen ST. Celestial Rotation: Its Importance in the Development of Migratory Orientation. Science. 1970 Dec 11;170(3963):1198–201. doi:10.1126/science.170.3963.1198

48. Alerstam T. Ecological causes and consequences of bird orientation. Experientia. 1990 Apr 1;46(4):405–15. doi:10.1007/BF01952174

49. Sokolovskis K, Lundberg M, Åkesson S, Willemoes M, Zhao T, Caballero-Lopez V, et al. Migration direction in a songbird explained by two loci. Nat Commun. 2023 Jan 11;14(1):165. doi:10.1038/s41467-023-35788-7

50. Sanchez-Donoso I, Ravagni S, Rodríguez-Teijeiro JD, Christmas MJ, Huang Y, Maldonado-Linares A, et al. Massive genome inversion drives coexistence of divergent morphs in common quails. Curr Biol. 2022 Jan 24;32(2):462–469.e6. doi:10.1016/j.cub.2021.11.019 PubMed PMID: 34847353.

51. Weissensteiner MH, Delmore K, Peona V, Lugo Ramos JS, Arnaud G, Blas J, et al. Combining Individual-Based Radio-Tracking With Whole-Genome Sequencing Data Reveals Candidate for Genetic Basis of Partial Migration in a Songbird. Ecol Evol. 2025;15(1):e70800. doi:10.1002/ece3.70800

52. Delmore K, Illera JC, Pérez-Tris J, Segelbacher G, Lugo Ramos JS, Durieux G, et al. The evolutionary history and genomics of European blackcap migration. Scordato E, Wittkopp PJ, editors. eLife. 2020 Apr 21;9:e54462. doi:10.7554/eLife.54462

53. Berthold P, Helbig AJ, Mohr G, Querner U. Rapid microevolution of migratory behaviour in a wild bird species. Nature. 1992 Dec;360(6405):668–70. doi:10.1038/360668a0

54. Berthold P, Terrill SB. Migratory behaviour and population growth of Blackcaps wintering in Britain and Ireland: Some hypotheses. Ringing Migr. 1988 Dec;9(3):153–9. doi:10.1080/03078698.1988.9673939

55. Delmore KE, Van Doren BM, Conway GJ, Curk T, Garrido-Garduño T, Germain RR, et al. Individual variability and versatility in an eco-evolutionary model of avian migration. Proc R Soc B Biol Sci. 2020 Nov 4;287(1938):20201339. doi:10.1098/rspb.2020.1339

56. Wynn J, Fandos G, Delmore K, Van Doren BM, Fransson T, Liedvogel M. Could bi-axial orientation explain range expansion in a migratory songbird? J Avian Biol. 2025;2025(1):e03196. doi:10.1111/jav.03196

57. Perea J. Ringing recovery of Yellow-browed Warbler in Andalucía confirms overwintering in consecutive winters. Br Birds. 2019;112:683–7.

58. Rogacheva H. The birds of Central Siberia. Husum; 1992.

59. Burrell AL, Sun Q, Baxter R, Kukavskaya EA, Zhila S, Shestakova T, et al. Climate change, fire return intervals and the growing risk of permanent forest loss in boreal Eurasia. Sci Total Environ. 2022 Jul 20;831:154885. doi:10.1016/j.scitotenv.2022.154885

